# Sap flow through petioles and petiolules reveals leaf-level responses to light and vapor pressure deficit in the tropical tree *Tabebuia rosea* (Bignoniaceae)

**DOI:** 10.1101/000711

**Authors:** Adam B. Roddy, Klaus Winter, Todd E. Dawson

## Abstract

Continuous measurements of sap flow have been widely used to measure water flux through tree stems and branches. However, these measurements lack the resolution necessary for determining fine-scale, leaf-level responses to environmental variables. We used the heat ratio method to measure sap flow rates through leaf petioles and leaflet petiolules of saplings of the tropical tree *Tabebuia rosea* (Bignoniaceae) to determine how leaf and leaflet sap flow responds to variation in light and vapor pressure deficit (VPD). We found that in the morning sap flow rates to east-facing leaves increased 26 minutes before adjacent west-facing leaves. Although leaves had higher integrated sap flow than their largest leaflet, this difference was not proportional to the difference in leaf area, which could be due to lower conduit area in petiolules than in petioles. In contrast to measurements on main stems, integrated daily sap flow was negatively correlated with daily mean VPD. Furthermore, leaves exhibited previously undescribed patterns of hysteresis in the sap flow-VPD and sap flow-PAR relationships. When hysteresis in the sap flow-PAR relationship was clockwise, the sap flow-VPD relationship was also clockwise; however, when hysteresis in the sap flow-PAR relationship was counterclockwise, the sap flow-VPD relationship displayed an intersected loop. These pattern differences highlight how substantially leaf-level processes may vary within a canopy and how leaf-level processes may not scale predictably to the stem level.

## INTRODUCTION

Approximately 90% of all water converted from the liquid phase to the vapor phase in terrestrial ecosystems moves through plants (Jasechko et al. 2013); in the tropics this amounts to an estimated 32×1015 kg of water per year (Hetherington and Woodward 2003). Almost all of this water transits through plant leaves. Understanding how leaves respond to abiotic drivers is important for modeling efforts at scales from leaves to landscapes (Jarvis and McNaughton 1986). At the leaf level, knowing which drivers impact transpiration most under daily and seasonally varying conditions is critical to understanding what may limit the distribution and abundance of species across the globe.

Various sap flow methods are commonly used to estimate almost continuously tree responses to environmental conditions for extended time periods (Marshall 1958, Granier 1985, Burgess et al. 2001, Vandegehuchte and Steppe 2012). These measurements are most often made on boles or large branches of trees and can be used to estimate canopy-level responses to changing environmental conditions (Oren et al. 1999a, Ewers and Oren 2000, Traver et al. 2010). Despite recent technical advancements in using sap flow measurements in stems to estimate canopy processes, a number of problems remain. First, resistance and capacitance in the hydraulic pathway creates time lags in water movement between different points in the hydraulic pathway (Köcher et al. 2013). For tropical trees, lags in sap flow between branches in the canopy and the stem base can approach an hour (Meinzer et al. 2004). Second, different parts of the canopy respond to environmental conditions largely independently (Brooks et al. 2003), such that sap flow measurements on branches or boles may provide only an average response of many leaves or branches. For example, east-facing branches of *Sequoiadendron giganteum* reached their maximum daily sap flow rates 6 hours before west-facing branches at the same height (Burgess and Dawson 2008). Although sap flow measurements on boles and branches have provided useful estimates of whole-tree water use (Wullschleger et al. 1998), their utility for describing leaf-level processes can be limited by a variety of factors including time lags, capacitance, hydraulic resistance, and variation in these factors along the root-to-leaf continuum.

Rarely have researchers attempted to measure sap flow rates through petioles of individual leaves (Sheriff 1972). Recently, Clearwater (2009) adapted the heat ratio method (HRM; Burgess et al. 2001) originally used for measuring sap flow through large stems to measure sap flow through small diameter stems, fruit pedicels, and leaf petioles. Slight variations of this method have proven useful in measuring sap flux through petioles under field conditions in the neotropics (Roddy and Dawson 2012, 2013, Goldsmith et al. 2013) and through stems of anatomically and phylogenetically diverse species of the South African fynbos fora (Skelton et al. 2013). These studies show that measuring sap flow directly adjacent to transpiring leaves can deepen our understanding of how leaves respond to variation in environmental conditions across a range of timescales. Placing sensors in close proximity to the transpiring leaves has the advantage of fine-scale measurements akin to leaf gas exchange without the disadvantage of enclosing leaves in a cuvette that removes the leaf boundary layer and otherwise modifies the leaf microenvironment.

Variation in sap flux is influenced by a variety of environmental conditions, including soil water availability, vapor pressure deficit (VPD), and solar radiation, and diurnal patterns may also vary seasonally (e.g. O’Grady et al. 1999, 2008, Zeppel et al. 2004). Over diurnal cycles, a change in an environmental variable in the morning does not always produce an equivalent response in sap flow as it does in the afternoon. Such a pattern is termed hysteresis and has been commonly observed in the sap flow responses to light and VPD. For example, at a given VPD that occurs both in the morning and again in the afternoon, sap velocity is higher in the morning (when VPD is increasing) than in the afternoon (when VPD is decreasing), creating a clockwise pattern of hysteresis throughout the day (Meinzer et al. 1997, O’Grady et al. 1999, Zeppel et al. 2004) that is consistent with hysteresis in canopy gas exchange (Takagi et al. 1998). Despite the ubiquity of hysteresis in sap flow data, its causes are rarely discussed. Hysteresis in a relationship indicates that factors other than the primary descriptor variable are constraining the response variable. For the relationship between sap flow and VPD, it is thought that hysteresis results from variation in hydraulic capacitance, resistance, or stomatal sensitivity to VPD (O’Grady et al. 1999). In the simplest case, no hysteresis in the sap flow-VPD relationship would mean that an increase and a decrease in VPD produce equivalent responses in sap flow and that there are no other factors influencing the sap flow response to VPD. However, a number of factors could cause deviation in the morning and the afternoon from this scenario of no hysteresis. First, trees often supply their morning transpiration from capacitive stores, which could elevate morning sap flow above that observed if there were no capacitance (Cowan 1972, Goldstein et al. 1998, Meinzer et al. 2004, 2008). Second, as stem water potential declines throughout the day, resistance in the hydraulic pathway increases, as is commonly observed in vulnerability curves (Meinzer et al. 2009), which could depress afternoon sap flow below that if there were no resistance. Both of these hydraulic factors, resistance and capacitance, could jointly be responsible for causing hysteresis in the sap flow-VPD relationship. Morning transpiration may draw largely on stored water, and as this hydraulic capacitor discharges, resistance may become important in depressing afternoon sap flow.

In contrast to VPD hysteresis, causes of hysteresis in the sap flow-light relationship are less clear. While the sap flow-VPD relationship commonly exhibits a clockwise pattern of hysteresis, the sap flow-light relationship generally exhibits a counterclockwise pattern (Meinzer et al. 1997, Zeppel et al. 2004). Zeppel et al. (2004) argued that counterclockwise hysteresis in the sap flow-light relationship results from the combination of (1) the difference in timing between peak light and peak VPD and (2) different stomatal responses to light and to VPD. Because VPD reaches its daily peak a few hours after light reaches its daily maximum at solar noon, sap velocity will be higher in the afternoon, when VPD is higher and stomata are fully open. Zeppel et al. (2004) argue that stomatal conductance saturates at relatively low light levels in the morning, and that above this saturating light level, VPD becomes the predominant driver of transpiration and sap flow. These responses probably vary between leaves acclimated to different microenvironments (e.g. between sun-and shade-leaves).

Using sap flow measurements on main stems of canopy trees to test these hypotheses for the causes of hysteresis are thus fraught with potential problems that focusing on leaves may circumvent. For leaves, sap flow responses to VPD are often similar to those for stems, although under some conditions leaves show different patterns (Roddy and Dawson 2013). In addition to clockwise hysteresis in the responses to VPD, sap flow through petioles sometimes exhibits an intersected loop (or ‘figure-eight’) pattern in response to diurnal variation in VPD. Determining the sap flow responses to environmental variables of individual leaves provides an opportunity to better elucidate important dynamics of plant water use. Furthermore, incorporating explicit measurements of sap flow to individual leaves could help to improve upon methods for scaling up to whole canopy processes.

In the present study, we measured sap flow rates through petioles and petiolules of saplings of the tropical tree *Tabebuia rosea* (Bignoniaceae) to understand how sap flow responds to variation in light and VPD. Because the figure-eight pattern of hysteresis in the sap flow-VPD relationship reported by Roddy and Dawson (2013) may result from an interaction with light, we also measured photosynthetically active radiation (PAR) levels on each leaf or leaflet to determine the conditions under which different patterns of hysteresis may occur. We were particularly interested in examining the differences in sap flow patterns between adjacent leaves and between leaves and leaflets because different microenvironmental conditions may cause sap flow patterns to differ between leaves on the same stem. Furthermore, differences in sap flux through petioles and petiolules may reflect variation in hydraulic architecture. If the hydraulic pathway constricts downstream, then sap velocity must increase as it moves towards leaflets. Our results highlight how measuring sap flow rates to individual leaves could deepen our understanding of the linkages between hydraulic architecture and plant water use.

## METHODS

### Plant Material

*Tabebuia rosea* (Bignoniaceae) grows to become a canopy tree in the lowland forests of central Panama. While adults are often deciduous, seedlings are evergreen with five palmate leaflets of varying size encircling the petiole. Plants were grown from seed in 20-liter, insulated pots outdoors under a glass roof at the plant growth facilities of the Smithsonian Tropical Research Institute in Gamboa, Panama, until a few days before sap flow measurements were begun. When the tops of the plants were ∼70 cm above the soil surface, they were transferred to a glass chamber to protect them from strong afternoon winds. The lower ∼1 m of this chamber was made of cement painted white, and doors on east- and west-facing sides of the chamber were left ajar to allow air circulation. At the beginning of the sap flow measurements, the tops of the plants were even with the top of the cement wall at the base of the chamber, and during the course of the experiment two new sets of leaves were produced. Pots were kept well-watered except for one week when water was withheld to determine how sap flow rates would respond to declining soil water. This week coincided with a dramatic increase in VPD. Sensors were installed when plants were approximately eight months old, a few days after transferring them to the glass chamber.

### Sap flow measurements

On each measured leaf, sap flow sensors were installed on the leaf petiole and on the petiolule of the middle, largest leaflet. On each plant, two adjacent leaves on opposite sides of the plant were chosen for measurement. At the time of installation, these leaves were the newest, fully-expanded leaves on each plant. Plants were positioned so that the axis defined by the two measured leaves on each plant were oriented east-west. Of the total 12 sensors installed, five failed, leaving two sensors on petiolules and five sensors on petioles.

Sap flow sensors and measurements were based on the design and theory of Clearwater (2009) with some slight modifications described previously (Roddy and Dawson 2012, 2013) and again briefly here. Sensors were constructed from a silicone backing and were connected to 10 cm leads with Molex connectors that were then connected by 10 m leads to an AM16/32 multiplexer and CR23X datalogger (Campbell Scientific Inc., Logan, UT). Sensors were held in place with parafilm, and sensors and connections were insulated with multiple layers of bubblewrap and aluminum foil at least 2 cm above and below the sensor. Consistent with previous applications of the HRM, we measured initial temperatures for 10 seconds prior to firing a 4-second heat pulse, monitored temperatures every 2 seconds for 200 seconds after the heat pulse, and initiated the measurement routine every 10 minutes.

The heat pulse velocity, *v_h_* (cm s^−1^), was calculated from the temperature ratio as:

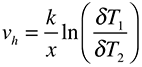

where *v_h_* is the heat pulse velocity in cm s^−1^, *k* is the thermal diffusivity (cm^2^ s^−1^), *x* is the distance from the heater to each of the thermocouples (cm), and *δT*_1_ and *δT*_2_ are the temperature rises (°C) above and below the heater, respectively (Marshall 1958, Burgess et al. 2001, Clearwater et al. 2009). We estimated the thermal diffusivity as:

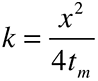

where *t_m_* is the time (seconds) between the heat pulse and the maximum temperature rise recorded *x* cm above or below the heater under conditions of zero sap flow (Clearwater et al. 2009). We measured *t_m_* every morning before dawn when atmospheric vapor pressures were lowest (between 0500 and 0600 hrs). At this time, the vapor pressure deficit was almost always below 0.3 kPa, and therefore we assumed *v_h_* was approximately zero. Thermal diffusivity, *k*, was calculated for each thermocouple (upstream and downstream) from these predawn measurements of *t_m_*, averaged for each sensor, and used to calculate *v_h_* from the heat ratios for the subsequent 24 hours. Measurements of *k* on nights with VPD always above 0.3 kPa were discarded and replaced with the most recently measured *k* when VPD < 0.3 kPa. We estimated the temperature ratio under zero-flow conditions by excising petioles and petiolules above and below the sensor at predawn at the end of the experiment, greasing the cut ends, placing the segments in a darkened box, and recording the temperature ratios for the subsequent ∼4 hours. The average of these zero-flow temperature ratios corresponded very well with the temperature ratios recorded predawn under low VPD (less than ∼0.3 kPa) conditions. The sensor-specific average temperature ratio under zero-flow conditions was subtracted from all calculated heat ratios. This corrected heat ratio was then used to calculate *v_h_*.

### Measurements of light and vapor pressure deficit

Light measurements were made using S1787 photodiodes (Hamamatsu Photonics, Hamamatsu City, Japan). Photodiodes were connected to 15 cm long copper wires with Molex connectors and then to 10 m leads, which were connected to a CR5000 datalogger measuring in differential mode. Circuits created by each photodiode were closed with a 100 Ohm resistor. Photodiodes were installed just above each leaflet with a sap flow sensor, and the photodiode was positioned to be parallel to the axis of the central vein of the leaflet. Light measurements were made every minute and averaged every 10 minutes. Voltage measurements from the photodiodes were converted to PAR based on a calibration of all photodiodes against a PAR sensor (LI-SB190, LiCor Biosciences, Logan, UT), during which time all sensors were situated adjacent to each other in a clearing that received full sunlight.

Vapor pressure deficit was calculated from temperature and relative humidity measurements made every 10 minutes with a HOBO U23 datalogger (Onset Computer Corp., Bourne, MA) that was housed in a covered, white, PVC, Y-shaped tube and hung level with the tops of the plants.

### Data analysis

All analyses were performed using R (R Core Team 2012). Raw velocity measurements were processed following previously published methods (Roddy and Dawson 2012, 2013, Skelton et al. 2013). Measurements of *v*_*h*_ were smoothed using the ‘loess’ function, which fts a polynomial to a subset of the data in a moving window of 35 points.

For analyses of structure (leaf vs. leaflet) or aspect (east-vs. west-facing leaves), sap flow measurements from individual sensors in each group were averaged. To estimate the total sap flow during the day and night, we integrated the time course of vh measurements for each day and night using the ‘auc’ function in the package *MESS*, which calculates the area under the curve using the trapezoid rule. Daytime was defined as being between 600 and 1800 hours, which corresponded to morning and evening twilight. To minimize the effects of nocturnal refilling, we defined nighttime as being between 100 and 600 hours, which assumed that diurnal water potential declines had mostly recovered within seven hours after sunset. To analyze the effects of VPD on integrated sap flow rates, we linearly regressed integrated sap flow against mean VPD. In all regressions for leaves and leaflets in the day and in the night, the linear model was determined to be as good or better than both the logarithmic and power functions by comparing the residual standard errors.

Differences in the timing of morning sap flow between east- and west-facing leaves were compared at a critical *v_h_* of 1.5 cm hr^−1^. We chose this critical value because it was higher than any measured nighttime velocities and lower than most daytime velocities. We estimated the time at which *v_h_* = 1.5 cm hr^−1^ by assuming a linear relationship (y = mx + b) between the two sequential morning measurements that spanned *v_h_* of 1.5 cm hr^−1^. To compare east-versus west-facing leaves and leaves versus leaflets, we used linear mixed effect models with day as the random variable, which accounts for repeated measures.

## RESULTS

Daily maximum VPD varied from 2.8 kPa to 6.9 kPa during the experiment, while daily maximum PAR at the top of the canopy varied from 1125 to 1640 *μ*mol m^−2^ sec^−1^. The daily maximum sap flow rate through petioles varied between 1.4 cm hr^−1^ to 4.5 cm hr^−1^. This lowest daily maximum *v*_*h*_ occurred at the end of a week without water, during which time five of the seven hottest, driest days occurred. Nighttime *v_h_* through petioles varied throughout the study, but was always below 1.0 cm hr^−1^ and below 0.5 cm hr^−1^ on all but seven nights. Overall, thermal diffusivity, *k*, ranged from 0.00136 to 0.00170 cm^2^ s^−1^. There were slight differences in *k* between sensors, but *k* was relatively constant throughout the experiment for each sensor, consistent with previously reported values for *k* from a diverse set of plant structures and species (Clearwater et al. 2009, Roddy and Dawson 2012, 2013, Skelton et al. 2013).

On every morning, sap flow rates to east-facing leaves increased more quickly than did sap flow rates to west-facing leaves. East-facing leaves had sap flow rates of 1.5 cm hr^−1^ on average 26 minutes before sap flow rates to west-facing leaves reached the same threshold (t = 5.67, df = 23, P < 0.001; Figure 1). In addition, on 18 out of 25 days, west-facing leaves reached their daily peak sap flow rate later in the day than east-facing leaves. However, sap flow rates to east-facing leaves did not decline any earlier in the evening than west-facing leaves, and east-facing leaves generally had higher nighttime sap flow rates than west-facing leaves, perhaps indicative of greater refilling.

**Figure 1.**
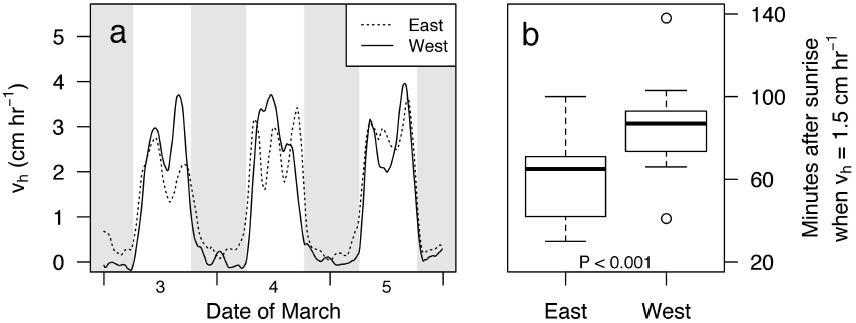
(a) Three days of sap flow for east-(dashed) and west-facing (solid) leaves. Tick marks on the horizontal axis indicate at midnight. (b) Boxplot of the time lag in sap flow rates between east-and west-facing leaves.

Patterns of sap flow to leaves and leaflets also differed. Leaflets generally had lower sap flow rates than leaves, and the daily integrated sap flow through petioles and petiolules differed significantly (t = 7.42, df = 24, P < 0.001; Figure 2). While water was withheld for one week, daily maximum sap velocities for both leaves and leaflets declined such that leaflet sap flow rates were about half of those to leaves (Figure 2a). On the day immediately following re-watering, leaves and leaflets had almost equivalent sap flow rates, which continued to increase on subsequent days despite declining daily maximum VPD during these days. Daytime integrated sap flow to leaves and leaflets was negatively correlated with mean VPD (Figure 3), both when including all days and when the last five days of the drought treatment were excluded (Table 1). There was a significant, negative relationship only between nighttime integrated sap flow of leaflets and mean nighttime VPD, but only when all data, including the drought days, were included. There was no relationship between nighttime VPD and integrated sap flow for leaves. Maximum *v*_*h*_ for leaves occurred at a slightly higher VPD than it did for leaflets (2.21 kPa vs. 2.06 kPa; grey symbols in Figure 3).

**Figure 2.**
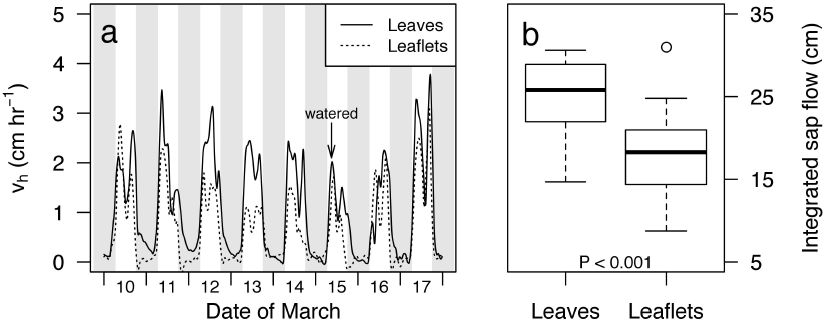
(a) Eight days of sap flow for leaves (solid) and leaflets (dashed). The first five days corresponded to the end of a week without watering. Tick marks on the horizontal axis indicate at midnight. (b) Boxplot of the daily time-integrated sap flow through leaves and leaflets.

**Figure 3.**
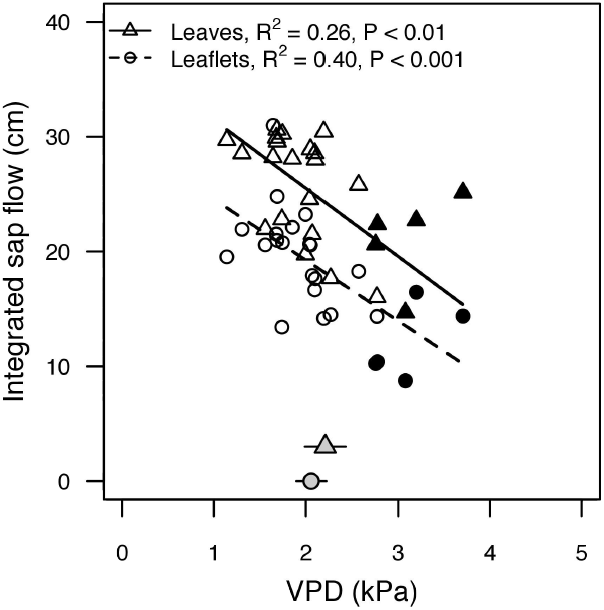
The relationship between daily integrated sap flow and mean daily VPD for leaves (triangles) and leaflets (circles). Black points represent the last five days during the drought treatment. Grey points at the bottom mark the mean VPD (and standard error) at which maximum daily sap flow occurred for leaves and leaflets.

**Table 1.** Summary statistics for the linear regressions between integrated sap distance and mean VPD for leaves and leaflets in the day and in the night, whether including data from the week of drought or not.

|  |  | Leaves |  |  |  | Leaflets |  |  |  |
| --- | --- | --- | --- | --- | --- | --- | --- | --- | --- |
|  |  | R<sup>2</sup> | t | d.f. | P | R<sup>2</sup> | t | d.f. | P |
| Day | All | 0.26 | 3.05 | 23 | **<0.01** | 0.40 | 4.15 | 23 | **<0.001** |
|  | No drought | 0.22 | 2.51 | 18 | **0.02** | 0.20 | 2.42 | 18 | **0.03** |
| Night | All | 0.04 | 0.94 | 23 | 0.36 | 0.28 | 3.19 | 23 | **<0.01** |
|  | No drought | 0.02 | 0.55 | 18 | 0.59 | 0.13 | 1.98 | 18 | 0.06 |

Patterns of sap flow hysteresis can be grouped into two classes, exemplified by data from two days from the same leaf shown in Figure 4. Data in Figure 4 are consistent with the patterns seen for other sensors on other days. The first type of hysteresis pattern is denoted by clockwise hysteresis in the relationship between *v*_*h*_ and VPD (Figure 4a). On this day, hysteresis in the *v*_*h*_-PAR relationship was also clockwise (Figure 4b). The second type of hysteresis is defined by an intersected loop, or figure-eight pattern, in the relationship between *v*_*h*_ and VPD (Figure 4d). On this day, the *v*_*h*_-PAR relationship had a counterclockwise pattern (Figure 4e). Nonetheless, the relationship between PAR and VPD for these two days was similar, showing a counterclockwise pattern for both days (Figure 4c,f).

**Figure 4.**
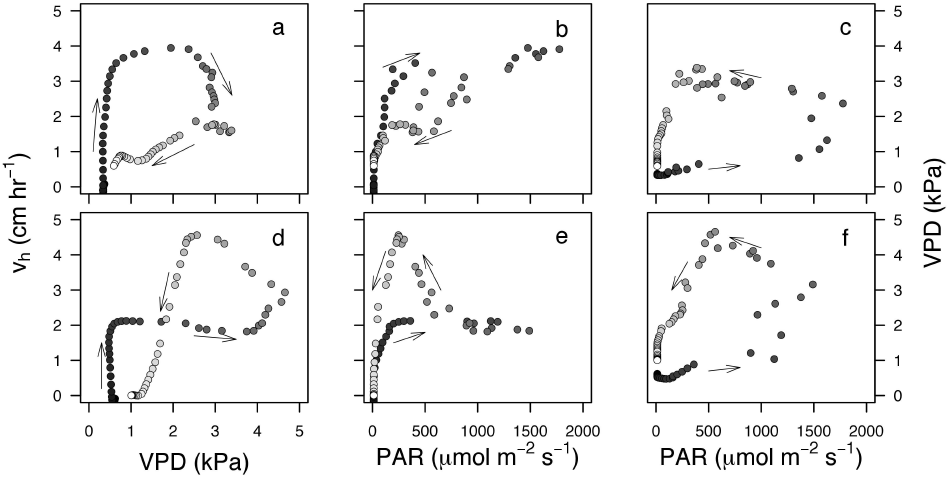
The pairwise relationships between *v_h_*, VPD, and PAR for two days (top and bottom rows). (a,d) The *v_h_*-VPD relationship differed between the two days, as did the (b,e) *v_h_*-PAR relationship. (c,f) However, the VPD-PAR relationship was approximately the same for the two days.

## DISCUSSION

Leaf physiology responds rapidly to changes in the leaf microenvironment in ways not previously appreciated (Zhang et al. 2013), and leaves may protect stems from low water potentials that can lead to loss of xylem functioning (Sperry 1986, Hao et al. 2008, Chen et al. 2009, 2010, Johnson et al. 2011, Bucci et al. 2012, Zhang et al. 2013). As a result, there has been burgeoning interest in the diurnal variability of leaf hydraulic functioning (Brodribb and Holbrook 2004, 2007, Johnson et al. 2009, Johnson et al. 2011, Wheeler et al. 2013). The new techniques for measuring leaf-level sap flow used here may become critical in quantifying diurnal variability in leaf functioning. Our results highlight how water use can differ significantly even between adjacent leaves, further justifying the need for fine-scale measurements like ours in quantifying leaf responses to environmental drivers. Notably, our sap flow measurements on individual leaves suggest that there may be important functional linkages between hydraulic architecture and water use. While a number of studies have shown these linkages for stems, substantially fewer attempts have been made to connect leaf hydraulic architecture to water use under natural conditions.

East- and west-facing leaves showed a number of differences in their patterns of daily sap flow (Figure 1a). Midday depression of *v*_*h*_ occurred in leaves on both sides of the plant, although not always at the same time of day, probably due to a combination of factors including differences in leaf energy balance and differences in the temporal and spatial dynamics of leaf water potential changes. Furthermore, aspect significantly affected the timing and rates of sap flow. Sap flow rates to east-facing leaves increased, on average, 26 minutes earlier in the morning than they did to west-facing leaves, due to earlier increases in leaf-level PAR to east-facing leaves than to west-facing leaves. Despite these differences in timing, leaf hydraulic conductance of east-and west-facing leaves may be similar if higher transpiration rates in east-facing leaves are accompanied by greater declines in leaf water potential. Leaf microclimatic conditions can cause substantial differences between even adjacent leaves on the same branch (Figure 1), which can influence the dynamics of branch sap flow (Burgess and Dawson 2008). How much influence water use by one leaf may have on water use by another, adjacent leaf is likely related to xylem hydraulic architecture. In addition to flowing longitudinally from roots to leaves, water may also flow laterally within a stem (MacKay and Weatherley 1973, James et al. 2003, Schulte and Costa 2010). The degree of this lateral flow varies among species and results from lateral connections between adjacent xylem vessels. Highly sectored xylem leads to close coupling of water uptake by roots on one side of the plant and water use by leaves on the same side of the plant. In this case, adjacent leaves on different sides of the plants would draw upon largely different pools of water in the stem. High sectoriality in xylem architecture allows plant parts to function independently, such that branches or leaves may compete little for water (Brooks et al. 2003, Orians et al. 2005). In contrast, highly integrated xylem (low sectoriality) leads to tighter hydraulic linkages between adjacent leaves on different sides of the stem axis. While we do not know how well integrated the xylem of adjacent leaves in *T. rosea* may be, orthostichous leaves (vertically aligned along the shoot axis) generally have more interconnected vasculature than do non-orthostichous leaves (those on different sides of a shoot; Watson and Casper 1984, Orians et al. 2005). Thus, adjacent east-and west-facing *T. rosea* leaves probably function more independently than would two east-facing, orthostichous leaves. Sap flow to individual leaves varies among leaves on the same branch, and the magnitude of this variation may itself vary among species depending on xylem architecture.

Patterns of sap flow through petioles were similar to patterns observed for petioles of other tropical species (Roddy and Dawson 2012, 2013), but, in some cases, different from patterns observed for main stems. On some days, patterns of hysteresis in the *v_h_*-VPD relationship were similar to those seen for main stems of canopy trees (Meinzer et al. 1997, O’Grady et al. 1999, 2008, Zeppel et al. 2004; Figure 4a). In this type of hysteresis, *v_h_* was higher in the morning than in the afternoon for a given VPD, creating a clockwise loop in the relationship between *v_h_* and VPD. On days when the *v_h_*-VPD relationship showed a clockwise loop, the *v_h_*-PAR relationship also had a clockwise hysteresis loop (Figure 4b). In contrast, for main stems, a clockwise *v_h_*-VPD loop is normally accompanied by a counterclockwise *v_h_*-PAR loop (Zeppel et al. 2004). On days with this first type of hysteresis, *v_h_* was higher in the morning than in the afternoon, with maximum daily *v_h_* occurring closer in time to peak PAR than to peak VPD. On these days, PAR peaked early in the day, saturating stomatal conductance and leading to high *v_h_* even when VPD was moderate. However, we observed a second type of hysteresis, characterized by a counterclockwise loop in the *v_h_*-PAR relationship (Figure 4e) that, unlike for main stems, was accompanied by a markedly different relationship between *v_h_* and VPD. When the *v_h_*-PAR relationship exhibited counterclockwise hysteresis, the *v_h_*-VPD relationship was characterized by an intersected loop, or figure-eight (Figure 4d). Although this intersected loop has been reported previously for main stems (Meinzer et al. 1999, O’Grady et al. 1999, 2008), its meaning has not been fully discussed or understood. This pattern occurred when afternoon *v_h_* was higher than morning *v_h_*, causing maximum daily *v_h_* to occur closer in time to peak daily VPD than to peak daily PAR. If morning transpiration is low and does not result in substantial water potential declines, then *v_h_* may peak in the afternoon when VPD peaks, as occurred on the second day shown. Why morning sap flow on this day was so low remains unclear but may be due to low water potentials, which we did not measure. Regardless, this second type of hysteresis exhibiting an intersected loop requires (1) a bimodal peak in the daily *v_h_* pattern (i.e. midday depression of gas exchange, which commonly occurs in tropical species) and (2) maximum daily *v_h_* to occur in the afternoon. Because slight midday depression of *v_h_* occurred on both days shown in Figure 4, midday depression alone may not lead to the intersected loop hysteresis. The second day did, however, have a higher afternoon VPD both in absolute terms (maximum of 4.77 compared to 3.44 kPa) and relative to PAR (Figure 4c,f), which was probably partly responsible for increased afternoon transpiration.

The most probable cause underlying such different patterns of hysteresis for individual leaves and for main stems is likely to be a matter of scale. Sap flow through stems integrates the individual sap flow responses of many leaves in drastically different microclimates. One of the most obvious sources of within-canopy variation is between different parts of a plant canopy that undergo different diurnal patterns of incident PAR, yet measurements on main stems ignore most of this within-canopy variation. In the present study, leaf aspect influenced patterns of sap flow, and previous studies on branches have shown that aspect influences both absolute rates of sap flow and the timing of peak sap flow within the day (Steinberg et al. 1990, Akilan et al. 1994, Martin et al. 2001, Alarcón et al. 2003, Burgess and Dawson 2008; Figure 1). Time lags between daily peaks of sap flow for east- and west-facing branches of large trees would result in different patterns of hysteresis depending on branch aspect (Burgess and Dawson 2008), and these patterns for branches may be similar to the second type of hysteresis (the figure-eight) we report for individual leaves. By measuring incident PAR to each leaf, we attempted in the present study to account for some of the variation in leaf microclimate that influences sap flow. However, we still ignored some important factors, such as leaf temperature and its effects on leaf saturation vapor pressure and the vapor pressure gradient (VPG) driving transpiration. This may be an acceptable oversight because atmospheric humidity has a greater impact on stomatal conductance than does leaf temperature (Fredeen and Sage 1999, Mott and Peak 2010). In addition to microclimatic variation, leaves and stems differ in their hydraulic architecture, which could influence sap flow patterns and hysteresis. Leaf water balance changes rapidly as efflux and influx of water vary asynchronously on the timescale of seconds (Sheriff and Sinclair 1973, Sheriff 1974). Water balance of stems may not change as rapidly, however, because of the compensatory effects of having numerous parallel pathways for water entry and loss. Thus, transpiration and sap flow may vary over much shorter timescales for leaves than for stems. Examining and quantifying sap flow hysteresis may provide new insights into hydraulic functioning in response to various abiotic factors influencing transpiration (Zeppel et al. 2004, Pfautsch and Adams 2013, Roddy and Dawson 2013).

Integrated daily plant water use, as measured by sap flow, generally increases with increasing mean and maximum daily VPD for plants from a wide variety of habitats, including canopy trees and shrubs (e.g. Zeppel et al. 2004, Pfautsch and Adams 2013, Skelton et al. 2013). However, in our experiment integrated daily leaf water use decreased with increasing mean daily VPD (Figure 3), whether days of declining soil water content were included in the analysis or not (Table 1). There was a significant negative relationship between integrated nocturnal sap flow and VPD for leaflets, but not for leaves, although this relationship was driven by very low nighttime sap flow during the drought (Table 1). For both leaves and leaflets, the VPD at which maximum daily *v*_*h*_ occurred was remarkably well conserved across days (2.21 and 2.06 kPa, respectively) and was, interestingly, the same whether maximum *v*_*h*_ occurred in the morning or in the afternoon (Figure 4a,d). These patterns opposite to those seen in main stems may result from higher than normal VPDs during our experiment. Leaves of *T. rosea* saplings may rarely encounter such high daytime VPD under natural conditions, and stomatal sensitivity to VPD may be responsible for the negative relationship we observed (Oren et al. 1999b). At VPD above ∼2 kPa, instantaneous sap flow rates often declined, consistent with stomatal closure to regulate transpiration rate and leaf water potential. For *T. rosea* saplings, the VPD at maximum *v*_*h*_ was higher than the VPD at maximum *gs* of other species, perhaps because of the higher than normal VPD during our experiment and the time lag between reaching maximum *gs* and maximum *v*_*h*_ due to hydraulic resistance.

In response to declining soil water availability, daily maximum *v*_*h*_ declined for both leaves and leaflets despite increasing VPD during this time. Rewatering caused an immediate increase in leaflet *v*_*h*_, such that it almost equaled leaf *v*_*h*_ (Fig. 2). Nonetheless, integrated leaflet sap flow was, on average, 30% less than leaf sap flow. Assuming the ratio of leaflet area to conduit cross-sectional area (LA:SA ratio) is the same for all leaflets, then instantaneous and integrated leaflet sap flow, as a fraction of leaf sap flow, should be proportional to leaflet area. However, both instantaneous and integrated leaflet sap flow were higher than this prediction, probably because cross-sectional conduit area of petiolules is lower than that of petioles. This could result from a combination of conduit taper and differences in the number of conduits between ranks (McCulloh et al. 2009, 2010). Although we did not measure conduit dimensions, our results highlight the potential linkages between leaf hydraulic architecture and diurnal patterns of water use at different scales. As of yet, there has been remarkably little effort to connect xylem structure-function relationships to continuous, sap flow measurements of plant water use.

## CONCLUSIONS

Understanding leaf-level responses to abiotic conditions is critical for modeling plant responses to future climate change. In the present study, we found that leaves often exhibit sap flow responses to abiotic drivers that are notably different from responses of stems, for two main reasons: (1) stems integrate over many leaves, each with their own microclimate and (2) moving the sap flow sensor closer to the sites of transpiration removes the confounding influence of capacitance distal to most stem or branch sap flow sensors. Thus, sap flow measurements on main stems may not accurately describe leaf-level processes. Furthermore, we found significant variation in sap flow patterns between adjacent leaves that are related to differences in the leaf microenvironment. Differences in sap flow between leaves and leaflets are likely due to differences in hydraulic architecture that influence patterns of water use. Future measurements of sap flow through petioles could better elucidate the biotic and abiotic drivers of transpiration dynamics under natural growth conditions.

## ACKNOWLEDGMENTS

Alexander Cheesman, Jorge Aguilar, Anna Schürkmann, Kevin Tu, and Joseph Wright provided technical assistance, and Daniel Johnson provided useful comments on a previous version of the manuscript.

